## Supplementary figures and images for "Multiple ETS Factors Participate in the Transcriptional Control of *TERT* Mutant Promoter in Thyroid Cancers"

### Supp_FigS1

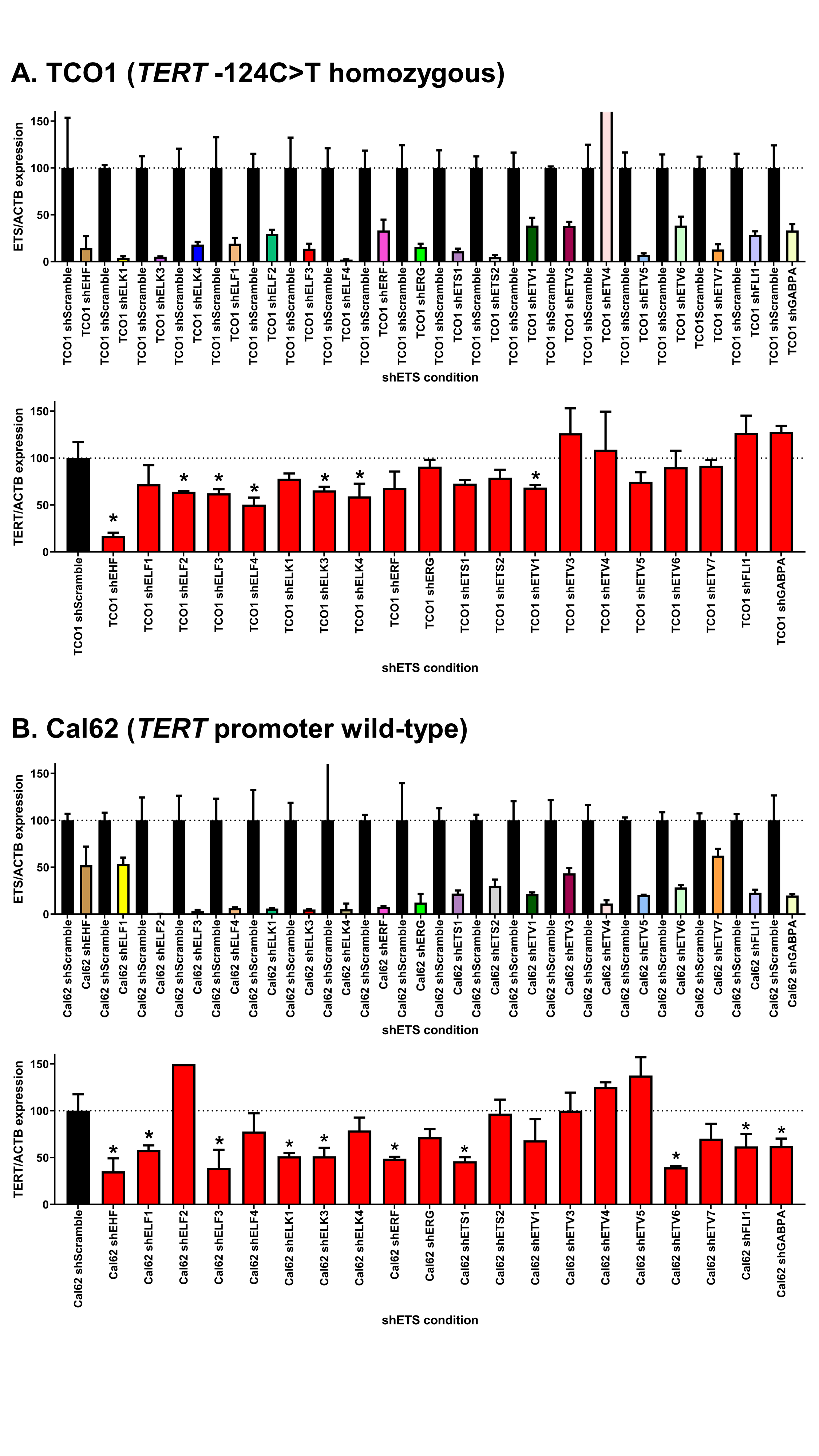

### Supp_FigS2

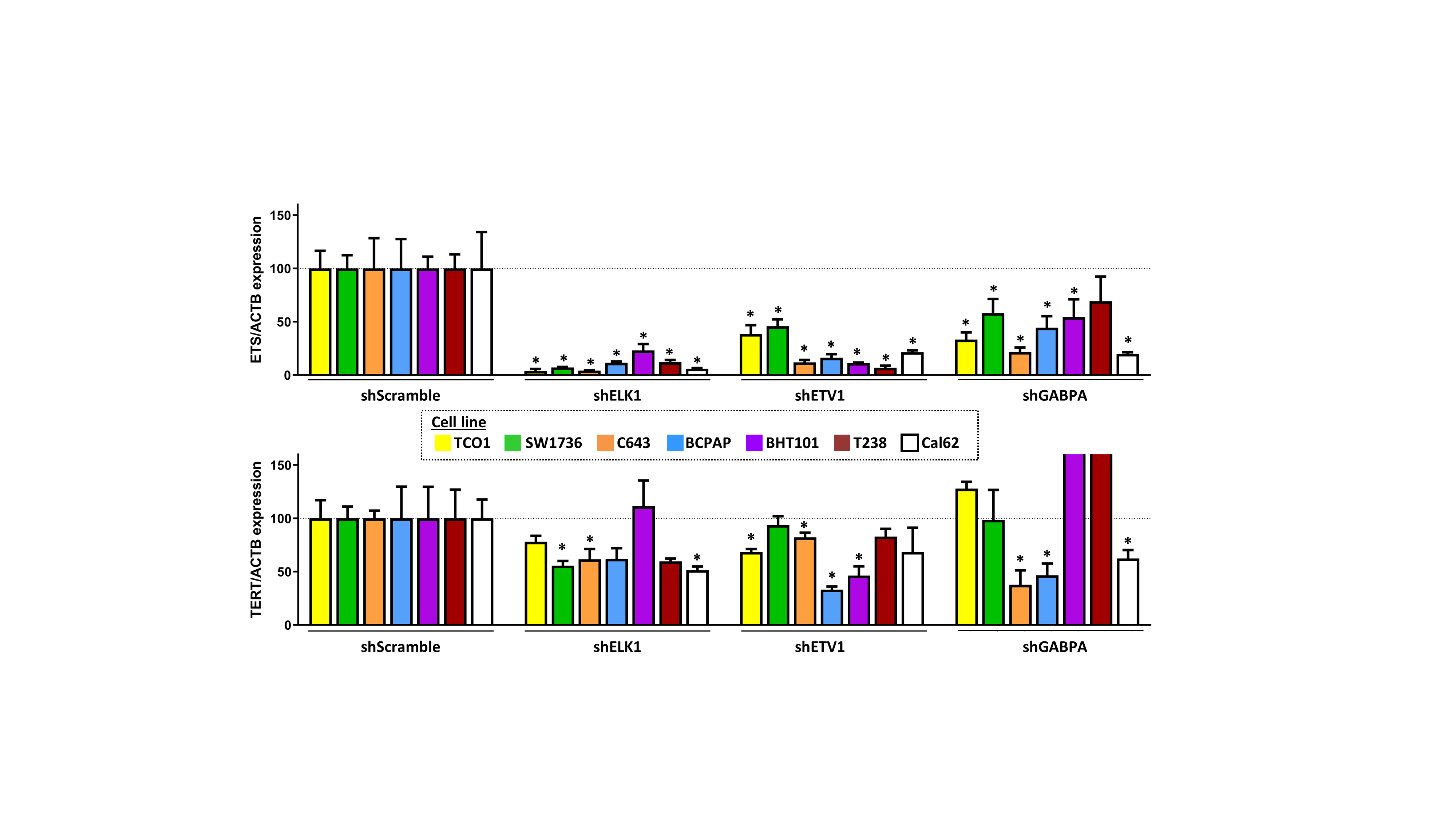
