## Supplementary material for "Multiple ETS Factors Participate in the Transcriptional Control of *TERT* Mutant Promoter in Thyroid Cancers": Supp_Figure_Legends

**Supplementary Figure Legends.**

**Supplementary Figure S1.** **Examples of ETS and TERT silencing assessment in thyroid cancer cell lines.** Expression levels of each of the 20 ETS factors upon shRNA-mediated silencing (top) panel and effects on TERT expression (bottom panel) in **A.** TCO1 cells (TERT -124C>T homozygous mutant); and **B.** Cal62 (TERT promoter wild-type). Expression levels for each gene are normalized to beta-actin (*ACTB*) and to their respective shScramble control. Asterisks indicate significant reduction of *TERT* levels for each of the shETS conditions compared to shScramble.

**Supplementary Figure S2.** **Results from the shRNA knockdown screen of ETS factors ELK1, ETV1 and GABPA in seven thyroid cancer cell lines.** Normalized expression levels for the indicated ETS factors (top panel) and TERT (bottom) are displayed. Asterisks indicate significant reduction of mRNA levels for each gene compared to its respective shScramble condition.
