## Supplementary material for "Multiple ETS Factors Participate in the Transcriptional Control of *TERT* Mutant Promoter in Thyroid Cancers": Supp_Tables

**Supplementary Tables.**

**Supplementary Table S1.** List of short hairpin RNA (shRNA) clones.

| **Clone ID (from GPP Web Portal)** | **Clone category** | **Target Sequence** | **Vector backbone** | **Original Target Gene ID** | **Original Target Gene Symbol** | **Match Regions** |
| --- | --- | --- | --- | --- | --- | --- |
| TRCN0000017078 | TRC 1.0 | CCGAGCTATGAGATATTACTA | pLKO.1 | 26298 | EHF | CDS |
| TRCN0000013856 | TRC 1.0 | CCTGCCGTAATTGTGGAACAT | pLKO.1 | 1997 | ELF1 | 5UTR, CDS |
| TRCN0000273855 | TRC 2.0 | GTCTCCTGAGTTAGGTATAAA | pLKO_TRC005 | 1998 | ELF2 | CDS |
| TRCN0000013863 | TRC 1.0 | CTCCCTAATTTATGTGCTATA | pLKO.1 | 1999 | ELF3 | 3UTR |
| TRCN0000013869 | TRC 1.0 | CGCGGAAGTCTTACTCAATAT | pLKO.1 | 2000 | ELF4 | CDS |
| TRCN0000007450 | TRC 1.0 | CCCAAGAGTAACTCTCATTAT | pLKO.1 | 2002 | ELK1 | 3UTR |
| TRCN0000329766 | TRC 2.0 | ACTCGTCCTTCACCATTAATT | pLKO_TRC005 | 2004 | ELK3 | CDS |
| TRCN0000013886 | TRC 1.0 | GCTTCTCTTACACCAGCATTT | pLKO.1 | 2005 | ELK4 | 3UTR, CDS |
| TRCN0000013909 | TRC 1.0 | CCTGCGCTATTACTATAACAA | pLKO.1 | 2077 | ERF | CDS |
| TRCN0000013914 | TRC 1.0 | GCTCATATCAAGGAAGCCTTA | pLKO.1 | 2078 | ERG | CDS |
| TRCN0000005591 | TRC 1.0 | CTGGAATTACTCACTGATAAA | pLKO.1 | 2113 | ETS1 | CDS |
| TRCN0000233984 | TRC 2.0 | GAGCAAGGCAAACCAGTTATT | pLKO_TRC005 | 23872 | Ets2 | CDS |
| TRCN0000013918 | TRC 1.0 | GCCGACTAAGAGAAGTTGTAA | pLKO.1 | 2114 | ETS2 | 3UTR |
| TRCN0000013923 | TRC 1.0 | GTGGGAGTAATCTAAACATTT | pLKO.1 | 2115 | ETV1 | 3UTR |
| TRCN0000013928 | TRC 1.0 | CCTCTTGACTTTCCTGGTTAT | pLKO.1 | 2117 | ETV3 | 3UTR |
| TRCN0000013934 | TRC 1.0 | GCAGAGCTTTAAGCAAGAATA | pLKO.1 | 2118 | ETV4 | CDS |
| TRCN0000273931 | TRC 2.0 | GAGCGATACGTCTACAAATTT | pLKO_TRC005 | 2119 | ETV5 | CDS |
| TRCN0000003852 | TRC 1.0 | AGGAGCTGGATGAACAAATAT | pLKO.1 | 2120 | ETV6 | CDS |
| TRCN0000016743 | TRC 1.0 | CCAGATGTGAAGCTCAAATTA | pLKO.1 | 51513 | ETV7 | 5UTR, CDS |
| TRCN0000005323 | TRC 1.0 | CCCATGAACTACAACAGCTAT | pLKO.1 | 2313 | FLI1 | CDS |
| TRCN0000304469 | TRC 2.0 | AGCTTAGTGTACAGGTAATTT | pLKO_TRC005 | 14390 | Gabpa | CDS |
| TRCN0000304508 | TRC 2.0 | ATGAACCAATAGGCAATTTAA | pLKO_TRC005 | 14390 | Gabpa | CDS |

Abbreviations: GPP= Genetic Perturbation Platform; CDS= coding DNA sequence; UTR= untranslated region

| **qPCR Primer List 5' -> 3'** | |  |
| --- | --- | --- |
| **Target Gene** | **Forward** | **Reverse** |
| *ETS1* | TACACAGGCAGTGGACCATC | CCCCGCTGTCCTGTGGATG |
| *ETS2* | CCCCTGTGGCTAACAGTTACA | AGGTAGCTTTTAAGGCTTGACTC |
| *ELF3* | CAACTATGGGGCCAAAAGAA | CAACCCTCAGTTCCGACTCT |
| *ELF1* | TGTCCAACAGAACGACCTAGT | GGCAGGAAAAATAGCTGGATCAC |
| *ELF2* | AAACTGTAGTGGAGGTGTCAACT | CATGGCTATCTGGTGATGTTGG |
| *ETV7* | TCAGCTGCTCCTTGATACCC | GCCCCGGTTCCTTCTTAAT |
| *ELF4* | CCTGATCTTTGAGTTCGCAAGC | AGTCCCGAGTACAGATGCAGT |
| *ETV1* | GGCCCCAGGCAGTTTTATGAT | GATCCTCGCCGTTGGTATGT |
| *ETV3* | GGTGGAGGGTATCAGTTTCCT | TGATGAATGGGTAGTTGGGCAT |
| *ETV4* | CAGTGCCTTTACTCCAGTGCC | CTCAGGAAATTCCGTTGCTCT |
| *ELK1* | TCCCTGCTTCCTACGCATACA | GCTGCCACTGGATGGAAACT |
| *ETV5* | CAGTCAACTTCAAGAGGCTTGG | TGCTCATGGCTACAAGACGAC |
| *ETV6* | CCCATTGGGAGAATAGCAGA | CAGGGCTCTGGACATTTTCT |
| *ERF* | ATTCATTGATGTGGGGTTGG | AGATGAAGAGCAGGCTGGTG |
| *EHF* | CCACCAGTCACCTTCCTGTT | GAGCCACTGCCTCTGATTTC |
| *ELK3* | ATCTGCTGGACCTCGAACGA | TTCTGCCCGATCACCTTCTTG |
| *ELK4* | ACTCAGCCGAGCCCTCAG | GGTGGCTTTTTGGAAGGTG |
| *GABPA* | AAGAACGCCTTGGGATACCCT | GTGAGGTCTATATCGGTCATGCT |
| *ERG* | TGGCTCAAGGAACTCTCCTG | ATAACTCTGCGCTCGTTCGT |
| *FLI1* | ATGACCACCAACGAGAGGAG | GTTCCTTGCCATCCATGTTC |
| *ACTB* | CTCTTCCAGCCTTCCTTCCT | AGCACTGTGTTGGCGTACAG |

**Supplementary Table S2.** List of primers used for quantitative PCR.
